# Microtubules Regulate Spatial Localization and Availability of Insulin Granules in Pancreatic Beta Cells

**DOI:** 10.1101/581330

**Authors:** Kai M. Bracey, Kung-Hsien Ho, Dmitry Yampolsky, Guoqiang Gu, Irina Kaverina, William R. Holmes

**Affiliations:** Department of Physics and Astronomy, Vanderbilt University; Department of Mathematics, Vanderbilt University; Quantitative Systems Biology Center, Vanderbilt University; Department of Cell and Developmental Biology, Vanderbilt University

**Keywords:** Glucose stimulated insulin secretion, computational modeling, membrane docking, anomalous diffusion

## Abstract

Two key prerequisites for glucose stimulated insulin secretion (GSIS) in Beta cells are the proximity of insulin granules to the plasma membrane and their anchoring or docking to the plasma membrane (PM). While recent evidence has indicated that both of these factors are altered in the context of diabetes, it is unclear what regulates localization of insulin and its interactions with the PM within single cells. Here we demonstrate that microtubule (MT) motor mediated transport dynamics have a critical role in regulating both factors. Super-resolution imaging shows that while the MT cytoskeleton resembles a random meshwork in the cells’ interior, MTs near the cells surface are preferentially aligned with the PM. Computational modeling demonstrates two consequences of this alignment. First, this structured MT network preferentially withdraws granules from the PM. Second, the binding and transport of insulin granules by MT motors prevents their stable anchoring to the PM. The MT cytoskeleton thus negatively regulates GSIS by both limiting the amount of insulin proximal to the PM and preventing/breaking interactions between the PM and the remaining nearby insulin. These results predict that altering MT structure in beta cells can be used to tune GSIS. Thus, our study points to a potential of an alternative therapeutic strategy for diabetes by targeting specific MT regulators.

## Introduction

Deregulated Glucose Stimulated Insulin Secretion (GSIS) results in diabetes, a disease that afflicts ~9% of the population in the USA (DeFronzo et al., 2015; Kahn et al., 2014; Stokes and Preston, 2017; Swisa et al., 2017). Thus, elucidating how GSIS is regulated is of fundamental importance in understanding glucose homeostasis at both the cellular and systemic level. Pancreatic islet beta cells are the insulin factories in the body. Here, insulin is produced / sorted through the ER and the Golgi (Fu et al., 2013), secretory insulin vesicles are generated at the TGN (Trans Golgi Network), and those vesicles mature into hard-core granules that are distributed through the cells cytoplasm for regulated secretion.

The major stimulant for insulin secretion is high glucose, whose entry into and subsequent metabolism in beta cells increases the ATP/ADP ratio, triggering insulin secretion (Rorsman and Ashcroft, 2018). The amount of secreted insulin is of a critical importance for metabolism and health, because over/under - secretion leads to hypo/hyperglycemia in patients. A main determinant of insulin secretion dosage at given stimuli is the number of readily releasable insulin vesicles, namely those that are biochemically capable of anchoring at the secretion sites and close enough to the plasma membrane to do so (Wang et al., 2009). Here we investigate how cells use the cytoskeleton to regulate this readily releasable pool (RRP) by controlling the number of granules near the plasma membrane as well as their availability for anchoring.

While numerous intra-cellular factors regulate the localization and availability of insulin granules, it has long been thought that the cytoskeleton has a critical role (Arous and Halban, 2015; Lacy, 1975; Roux et al., 2016). Cytoskeletal polymers microtubules (MTs) and MT-dependent molecular motors are the major transport system in mammalian cells (Barlan and Gelfand, 2017; Vale, 2003). In many cell types, MTs extend toward cell periphery in radial (mesenchymal cells) or parallel (neurons, columnar epithelia) arrays, allowing them to serve as long-distance transport highways, for example for delivery of secretory vesicles, among other functions (Baas and Lin, 2011; Kapitein and Hoogenraad, 2011; Muroyama and Lechler, 2017; Vinogradova et al., 2009). In pancreatic beta cells, MTs also serve for intracellular transport (Donelan et al., 2002; McDonald et al., 2009; Varadi et al., 2002), but MT function in secretion is complex and incompletely understood. In the long-term, MT depletion inhibits new insulin granule formation by interfering with insulin transport through the endoplasmic reticulum (ER) and the Golgi ((Malaisse-Lagae et al., 1979) and our unpublished data). A number of observations indicate that prolonged insulin secretion is attenuated in the absence of MTs (Boyd et al., 1982; Lacy et al., 1972), which could be explained by lack of new granule production/delivery (Hoboth et al., 2015). Moreover, without MTs, the net movement of existing secretory insulin granule movement, although not abolished, is significantly slowed (Zhu et al., 2015). Interestingly, in our recent finding short-term depletion of MTs resulted in immediate facilitation of exocytosis and, as a result, increased GSIS, which is consistent with earlier findings (Devis et al., 1974; Somers et al., 1974). Moreover, MT enrichment in beta cells both in taxol-treated islets and in diabetic mice (Zhu et al., 2015) was associated with decreased secretion. Thus, while all studies agree that MT-dependent transport is needed for new insulin granule production, it is not readily apparent how or why MTs regulate secretion of the RRP, or how transport of existing granules is linked to GSIS. Here we test the hypothesis that this link between MT transport and secretion is a consequence of the cytoplasmic architecture of beta cells.

One important feature of beta cell cytoplasm is the abundance of premade insulin granules in a resting cell. Estimates indicate individual beta cells contain on the order of 10,000 insulin granules (Dean, 1973), each 100-300 nm in diameter (Rorsman and Renstrom, 2003), which are tightly packed in the cytoplasm across a cell. At any stimulation, only a small portion of these vesicles was secreted [~1% within an hour of high glucose stimulation, (Rorsman and Renstrom, 2003)]. This raises the question, why these abundant vesicles should be transported for GSIS. Additionally, granule motions analyzed in beta cell culture models (Tabei et al., 2013) and in intact pancreatic islets (Zhu et al., 2015) were found to be random and undirected. This is not surprising given that super-resolution imaging of the MT cytoskeleton in intact islets has indicated that beta-cell MTs form a spaghetti-like random meshwork (Zhu et al., 2015), which is very different from directed MT arrays in cells that use MTs for directional long-distance transport. Thus, even if transport were important for GSIS, what would random transport on an unstructured MT meshwork accomplish and how would it influence GSIS, is unclear.

Our prior data provide a clue to how MTs influence GSIS. In the absence of MTs, high glucose stimulation leads to accumulation of granules at the cell periphery (Zhu et al., 2015), possibly due to the stimulation of glucose-dependent priming/docking (Gandasi et al., 2018). Interestingly, the presence of MTs prevents this excess accumulation, suggesting MT transport may regulate granule localization even when motions are random and undirected. TIRF microscopy data points to two possible mechanisms for this regulation. First, quantification of delivery and withdrawal of granules from the cell periphery demonstrates that MT-dependent transport is required to maintain the proper balance between delivery and removal (Zhu et al., 2015). Second, the motions of granules near the membrane (in the TIRF field, within ~200nm of the plasma membrane) are predominantly parallel to it (Varadi et al., 2002), indicating there may be structure to the MT network near the membrane and that motions may not be random there. To clarify which mechanism is likely supported by the beta cell MT network, we utilize super-resolution microscopy to image the structure of the MT meshwork near the plasma membrane and computational modeling to assess how the interactions between granules, the that meshwork, and the plasma membrane influence GSIS.

There are generally two populations of MTs in cells: dynamic MTs that are undergoing dynamic disassembly (Brouhard and Rice, 2018; Mitchison and Kirschner, 1984), and stable MTs, characterized by the presence of detyrosinated tubulin (Glu-tubulin) among other modifications (Garnham and Roll-Mecak, 2012; Hammond et al., 2008; Roll-Mecak, 2019). Glucose alters the MT network in potentially important ways While glucose only modestly alters the density of MTs in cells, it does make the network significantly more dynamic by both destabilizing and depolymerizing stable MTs and increasing the rate of new MT nucleation (Zhu et al., 2015) and MT growth rates (Heaslip et al., 2014). Glucose is also well known to activate docking molecules, which are necessary for GSIS. This body of work thus suggests that glucose stimulation influences granule transport, which in turn alters GSIS.

We have hypothesized that MTs have a dual role in negatively regulating GSIS: MTs 1) enhance withdrawal of granules from the periphery to the interior and 2) prevent anchoring and subsequent secretion of those at the periphery (e.g. by preventing the formation of or breaking bonds between granules and the anchoring machinery). While this is a compelling hypothesis, our understanding of MT control on cytoplasmic distribution of insulin granules remains fragmented and insufficient. In particular, the abundance of insulin, the apparently random nature of the MT network, and the seemingly random but complex nature of granule motions (Tabei et al., 2013) makes it difficult to deduce how MTs influence GSIS. To test this hypothesis, we will develop a computational model of intra-cellular insulin dynamics to investigate how MT dynamics influence insulin localization and availability. The basic elements of this model (e.g. transport rates and MT binding rates) will be calibrated to data. It will then be tested against independent results, including TIRF observations of peripheral granule densities along with quantification of GSIS under different conditions, to determine under what conditions the model matches observations. In this way, the model and experimental observations will be jointly used to infer how interactions between MT dynamics, granule dynamics, and membrane anchoring influence GSIS.

In this study, we investigate two specific questions, both of which are important to understanding GSIS. How does MT transport influence the density of granules near the plasma membrane and how does the binding of granules to the MT cytoskeleton influence their membrane anchoring, both of which are a pre-requisite to exocytosis. Given our focus on the dynamics of granules near the plasma membrane, we will quantify the structural characteristics of the MT network near the membrane (directionality in particular) in pancreatic beta cells. This data is used in conjunction with prior 3D tracking of granule motions (Zhu et al., 2015) to develop and simulate a discrete, two-dimensional computational model of insulin granule dynamics within a single cell. Results of this modeling supports the aforementioned hypothesis that MT transport negatively regulates GSIS in two important ways: by 1) increasing the rate of transport of granules away from the plasma membrane and 2) reducing the ability of those that are near the membrane to stably anchor to it.

## Results

### Peripheral MTs in islet beta cells are co-aligned with the cell border

Prior imaging of intact islets indicate (Zhu et al., 2015) that MT network in beta cells appears to lack previously assumed radial directionality characteristic commonly seen in mesenchymal cells in culture, and instead resembles an undirected random mesh. However, directionality of MTs in beta cells has not been quantitatively characterized, and functional consequences of variable directionality have not been computationally assessed. Here we analyze directionality of MT in beta cells using a custom image-analysis algorithm. In subsequent sections, we use computational modeling to assess the consequences the type of MT organization near the plasma membrane.

Intact mouse pancreatic islet were isolated and equilibrated according to a standard protocol. After a pretreatment in low and high glucose conditions, islets were fixed and immunostained for insulin to distinguish beta cells, e-cadherin for cell border identification, and for tubulin for MT network identification. Confocal stacks of whole-mount islets were deconvolved for increased resolution (Fig. 1 A, B). Single 2D slices of MT images were subjected to threshold (Fig. A,B, second from the right) and directionality of MTs was determined in respect to the cell border (Fig. A, B, right). Every pixel of the image was analyzed, while inconclusive pixels were disregarded. Subsequently, MT directionality was quantified as a function of the distance from the cell border (Fig. 1C,D).

Our results indicate that away from the cell border, in the cell interior, the MT network lacks directionality and resembles a random interlocked mesh (Fig. 1C,D, right). MTs within a narrow peripheral region however exhibit a significant co-alignment with the cell border (Fig. 1C,D, left). The fraction of MTs that were border-aligned was similar in low and high glucose, although the number of detectable pixels was lower in high glucose, consistent with our previous finding of partial MT destabilization under high glucose conditions (Zhu et al., 2015). Interestingly, visualization of long-lived (stable) MTs by detyrosinated (Glu-) tubulin staining detected many Glu-MTs co-aligned with the cell periphery in low (Fig. 1 E), but not in high (Fig. 1 F) glucose. Since MTs parallel to the cell border are still observed in high glucose (Fig.1 C), we conclude that stability of this peripheral bundle is significantly diminished by glucose-triggered MT destabilization.

**Figure 1:**
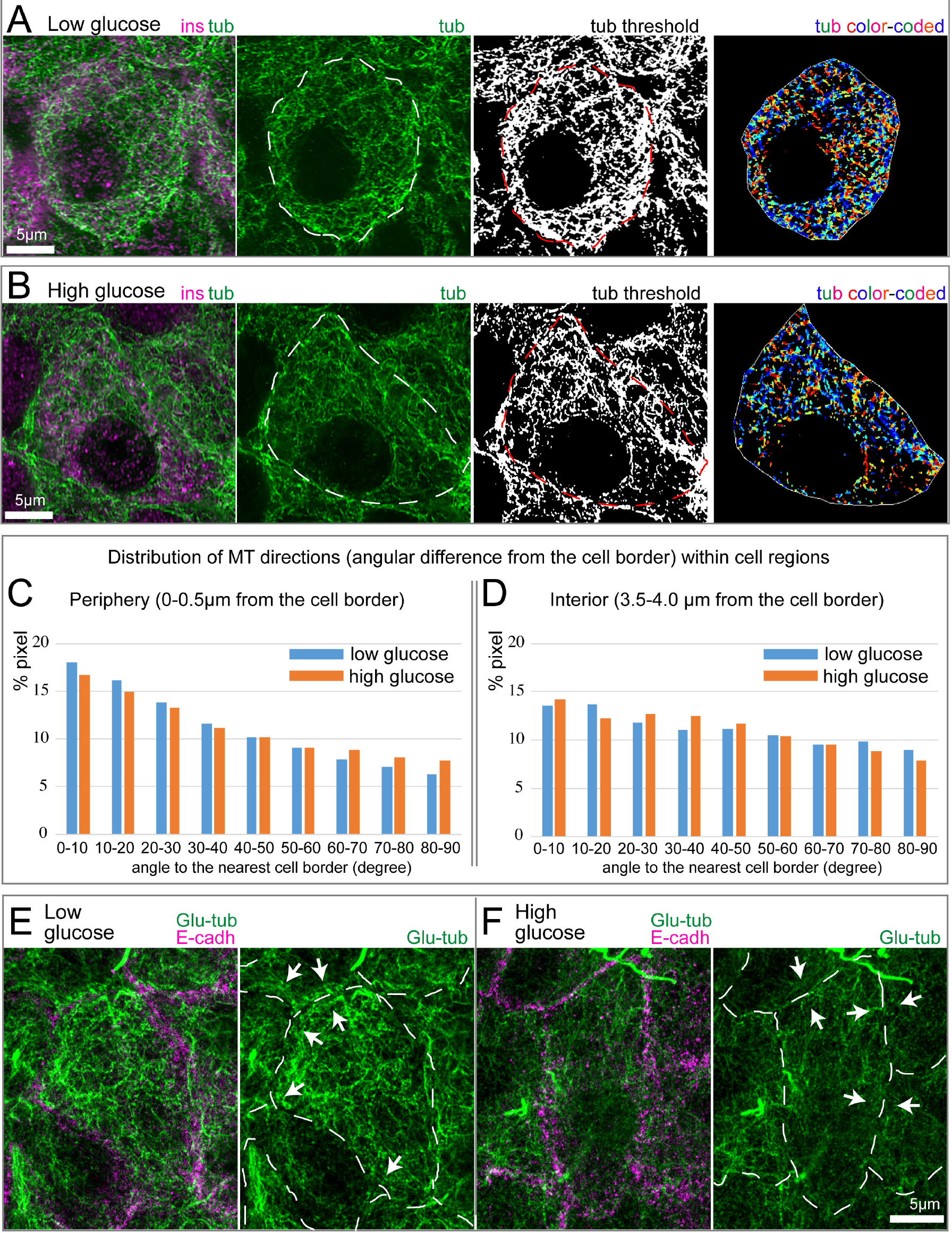
MTs parallel to the cell edge in beta cells are destabilized in high glucose. **Panels A, B:** Examples of MT directionality quantification in low (A) and high (B) glucose. Tubulin, green. Insulin (beta-cell marker), magenta. Cell outline, as detected by e-cadherin staining, is shown as dotted line on tubulin and thresholded images. The image on the right shows color-coded MT directions: parallel to the cell edge, blue; perpendicular to the cell edge, red. **Panels C, D:** Histograms of MT directionality within two cell regions: periphery (C) and interior (D). Percentage of tubulin-positive pixels in the analyzed cell population is shown. MTs at the periphery tend to be parallel to the edge. Low and high glucose do not differ. N=12 and 11 cells for low and high glucose, respectively. Pixel numbers in the analysis: 71759 (low, periphery); 9622 (low, interior); 43747 (high, periphery); 5833 (high, interior). Note that a lower number of pixels was identified in high glucose, consistent with the fact that MTs are destabilized under this condition. **Panels E, F:** Stable MT marker detyrosinated (Glu-) tubulin (green) in low (E) and high (F) glucose. Cell-cell adhesions are stained for E-cadherin (magenta) in left-hand panels and outlined (dashed line) in right-hand panels. Note multiple Glu-tubulin-positive MTs parallel to the cell border (arrows) in low glucose (E). Glu-tubulin content is decreased in high glucose (F), indicating MT destabilization both across the cell and at the cell border (arrows).

### Microtubules generate counter propagating density gradients that differentially deliver and remove granules from the cell periphery

In the remainder of this study, we develop and utilize computational modeling to assess how the MT organization feature described above, along with MT and non-MT-dependent transport processes influence insulin granule localization. For specific model and implementation details as well as a discussion of how parameters for the model were calibrated to data or chosen, see the Methods section. Briefly, the model used here is comprised of a discrete, 2D network of non-interacting MTs along with a population of granules that undergo MT dependent and MT independent motion. These granules are assumed to both bind and unbind to the MT network and to anchor to the plasma membrane when glucose is present.

Based on the above analysis, we consider a range of assumptions for how MTs interact with the plasma membrane. Computationally, we generate the MT network by essentially growing individual MTs from a random seed location. We consider two assumptions for how MTs interact when the reach the border: they either terminate or bend and grow parallel to the periphery. By varying the likelihood of each *in silico* MT doing one or the other, we can vary the net orientation of the resulting peripheral network from being highly aligned to having no alignment. As we do not know *a priori* the significance of this orientation on insulin granule dynamics, we explore the influence of this and the other aforementioned factors on peripheral granule density.

Quantification (Figs. 2 a-c) of the steady state number of granules (total, MT bound, and unbound) near the cell border as a function of both the total number of cellular MTs and the peripheral alignment of MT’s indicate both influence granule densities. Here, “peripheral alignment” of MTs refers the fraction of MTs that interact with the boundary that grow parallel to it.

**Figure 2:**
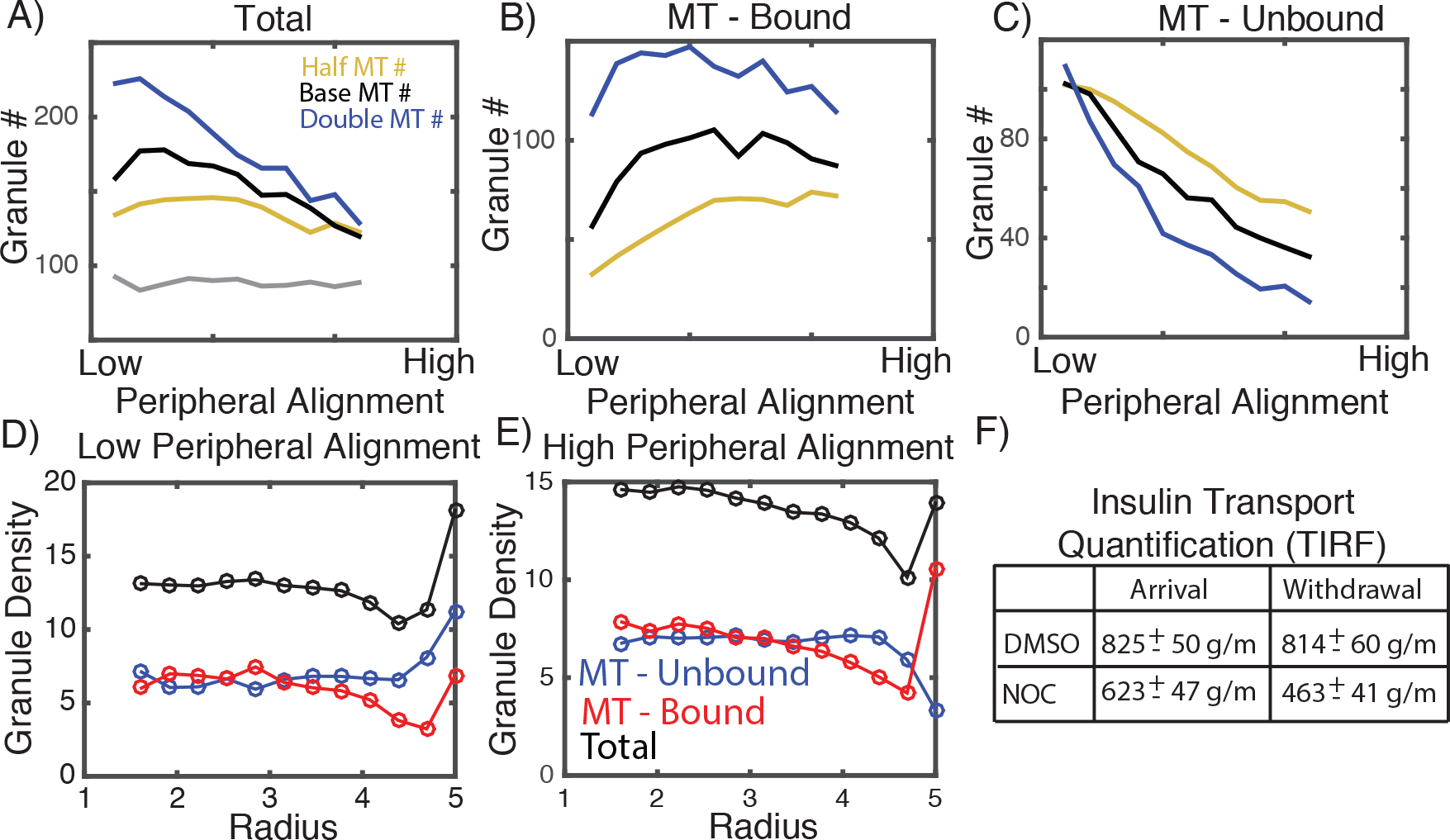
Effect of microtubules on peripheral localization of insulin granules. **Panel A:** Total number of granules located within 250 nm of the cell membrane as a function of the peripheral density of MTs as well as the total density of MTs. **Panel B:** Number of MT bound granules. **Panel C:** Number of unbound granules. **Panels D, E:** Radial distribution of bound, unbound, and total granules in low and high peripheral density cases. **Panel F:** Table depicting change in granule arrival delivery to the cell periphery in cells from experiments in our previous publication (data reproduced from (Zhu et al., 2015)).

Results (Fig. 2a) show the presence of MT’s always leads to an enrichment of granules near the cell periphery relative to densities when the MT network is completely removed. This is true for all MT densities and peripheral MT alignment conditions tested. In this study, we assumed that motors carrying granules stall at the tip of MT’s. To ensure this is not the source of these results, we carried out identical simulations where motors are assumed to disassociate at the tip (SM Fig. 1) and find similar results. Similar simulations were also performed where granule motions are purely diffusive rather than sub-diffusive, again with similar results (SM Fig. 1). This suggests that the MT network serves as a sponge of sorts that enriches granule densities near the cell border, which is likely the result of the increased density of MT tips near the cell periphery.

Interestingly, this enrichment effect appears to be weakened when the MT network is more aligned at the cell periphery. Specifically, when the MT network at the periphery is more aligned, fewer granules localize to the boundary. Further, when the network is highly aligned, peripheral density becomes essentially independent of total MT density. Further inspection of the peripheral densities of bound and unbound granules as a function of peripheral alignment (Figs. 2 b, c) provides a clue as to the cause of this observation. Increasing peripheral alignment has little influence on the number of bound peripheral granules but leads to a substantial reduction in the density of unbound peripheral granules. On net, this yields the observed inverse relationship between total peripheral granule density and peripheral MT alignment.

This suggest that an enrichment of peripherally oriented MTs would serve to 1) increase the total binding of peripheral granules to the MT network (thus reducing unbound granule numbers) and 2) transport those excess granules toward the cell interior (thus leaving the fraction of bound granules relatively unchanged). Critically, this transport of bound granules away from the periphery is not due to directional motions of kinesin or dynein motors since all granule motions are random and undirected. This raises an important question. If, at equilibrium, peripherally aligned MT’s serve to soak up and transport granules away from the periphery, what counter balances that net transport? To answer this, we quantified (in simulations at steady state) the density of bound and unbound granules as a function of radial distance from the cell center (Fig. 2d, e) for the two extreme cases of low and high alignment of peripheral MT’s. In the highly aligned case (Fig. 2e), bound and unbound granule densities exhibit opposing density gradients with unbound granules exhibiting a peripheral deficit and bound granules a peripheral enrichment. When the alignment of peripherally aligned MTs is low (Figure 2d), these opposing gradients are not present. Thus, when there is a substantial number of peripherally aligned MT’s, bound and unbound granules form counter-propagating gradients with unbound granules flowing from the interior to the periphery and bound granules flowing from the periphery to the interior.

This counter-propagating gradient theory is consistent with our prior observations. We found, using TIRF microscopy, that the application of NOC and glucose led to an ~25% reduction in granule delivery but an ~43% reduction in granule withdrawal (Zhu et al., 2015). Thus removal of granules was more substantially impacted by removal of MT’s than delivery, consistent with the counter-propagating gradient hypothesis where MT’s generate a net flow of granules from the periphery to the cell interior. In combination, these results suggest that the peripherally aligned network of MTs maintains a balance between delivery and withdrawal of granules and prevents excess accumulation near the plasma membrane.

### Changes in radial diffusion due to MT alignment are the source of these counter-propagating gradients

We investigate two potential effects of peripheral MT alignment on insulin localization. Enrichment of these peripherally aligned MTs could serve to either 1) increase binding of granules to MTs or 2) restrict the radial motility of bound granules. In the discrete model it is impossible to separate these effects; increased peripherally aligned density will necessarily influence both. To assess the relative importance of these in potentially generating the aforementioned counter-propagating gradients, we construct a simplified continuum model of granule dynamics where the two can be separately modulated.

For this continuum model, we consider concentrations of granules rather than individual granules and use partial differential equations (PDEs) to describe time evolution of the spatial concentrations. Since the presence of these counter-propagating gradients in the prior study was not the result of sub-diffusion (Fig. S2), we consider the motions of both bound and unbound granules to be purely diffusive. This greatly simplifies the model, allowing it to be described by standard reaction diffusion PDE’s. For simplicity, the cell is considered to be a radially symmetric circle and we model only the dynamics in the radial direction since steady state distributions in the discrete model depend on radius but not angular position in the cell. This reduces the model to a one dimensional, radially symmetric system that further simplifies calculations while allowing us to assess the influence of these factors on radial density.

This model encodes three essential components of the discrete model: 1) the ability of granules to bind and unbind from MT bound to unbound states, 2) diffusion of unbound granules, and 3) diffusion of bound granules. It does not however explicitly include discrete MTs. Rather, bound and unbound forms are assumed to move with different rates of diffusion (faster for bound). To determine how increases in the rate of MT binding and decreases in the rate of radial diffusion at the cell periphery (due to MT enrichment) influence distributions of bound and unbound granules, we define a 250nm zone near the cell border where MT binding rates (k_on_) and speed of bound granule radial diffusion (D_r_) are selectively modulated. The benefit of this continuum approach is that we can separately and selectively change these two parameters near the cell border to assess their influence in isolation.

The model was simulated for a range of different fold increases in the binding rate and fold decreases in the rate of bound granule radial diffusion (again, these parameters are modulated only near the periphery). Results (Fig. 3) show that changes in binding and radial diffusion rates have different roles in setting up these counter-propagating gradients. An increase in the binding rate is sufficient to induce a depletion of unbound peripheral granules, but not sufficient to induce a significant gradient in bound granules. A reduction in the rate of radial diffusion (in combination with the increase in binding rate) does however lead to a substantial enrichment of bound granules at the periphery. Thus, a moderate increase in the MT binding rate in combination with a substantial decrease in the rate of bound granule radial diffusion is required to explain the counter-propagating gradients seen in the discrete model. Both would be expected to occur if MTs are enriched at the cell periphery.

**Figure 3:**
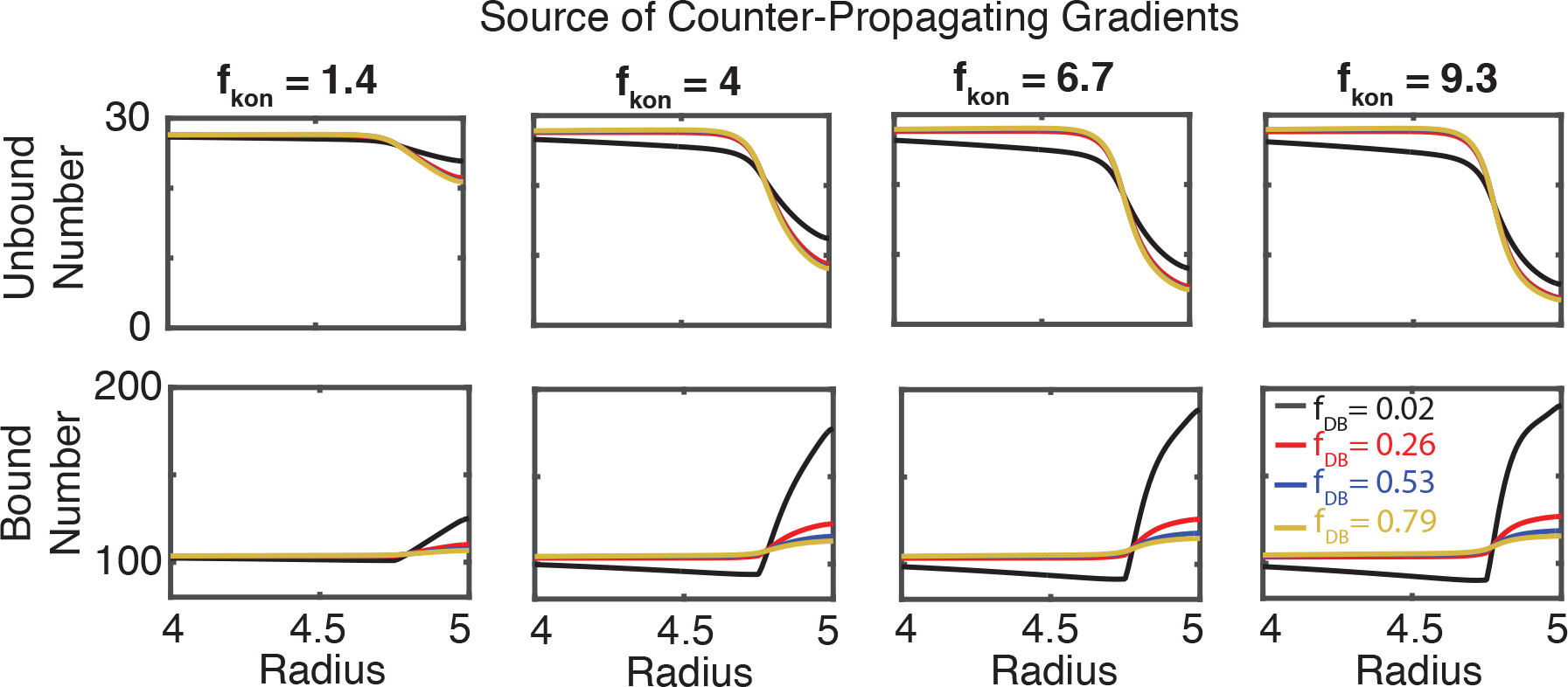
Effect of MT binding propensity and reduction in radial diffusivity on granule densities. Simulations of radially symmetric continuum model where the rate of MT binding is increased by a factor (f_kon_) and rate of radial diffusivity is decreased by a factor of (f_DB_) within 250nm of the cell membrane. Top (bottom) panels show the steady state insulin densities as a function of radius in different conditions. The notation f_DB_ = 0.02 and so forth indicates the base value of the relevant parameter is multiplied by the given value in the peripheral region (DB → 0.02*DB, in this case).

### The anomalous nature of granule motion alters localization of granules near the cell membrane in a MT-dependent fashion

It is well established that insulin granule motion (like the motion of many entities within the cell) is sub-diffusive (Zhu et al., 2015), characterized by mean squared displacement curves obeying *MSD* = *Dt*^*α*^ where “D” is the generalized diffusion coefficient and “α” is the diffusive exponent: α=1 corresponds to regular diffusion while α<1 indicates sub-diffusion. For insulin granules, it has previously been found that α~0.75 (Tabei et al., 2013), indicating significant anomalousness of diffusion. It has been further found insulin granule motions exhibit characteristics of fractional Brownian motion (Tabei et al., 2013), which is often associated with visco-elastic drag effects arising from the complex and crowded nature of the cells cytoplasm. While numerous studies have investigated the anomalous nature of random particle motion in cellular environments (see (Hofling and Franosch, 2013) for a comprehensive review), to our knowledge, the effect of visco-elastic sub-diffusion on the spatial distribution of particles at equilibrium (granules in this case) has not been investigated. Here we assess how this feature of motion, and its changes due to alterations in the MT cytoskeleton, affects steady state spatial densities of granules within the cell.

It is well established that when particles obey standard random / Brownian motion, spatial distributions of particles tend to homogenize within a spatial domain. To determine if this is the case when motions are more complex and governed by visco-elastic sub-diffusion, we simulated the spatial distribution of 1000 non-interacting granules over time for different values of the *D* and α parameters (Figure 4). In order to independently assess the influence of anomalous motions, we initially consider only the two-dimension motions of granules independent of MT’s. Results show that when granule motions are sub-diffusive, there is a significant depletion of granules at both the cellular and nuclear borders (Fig. 4d). Furthermore, as motions become more sub-diffusive (smaller α) or faster (larger *D*), this depletion near the cell border becomes more substantial (Figs. 4 a,d,f).

**Figure 4:**
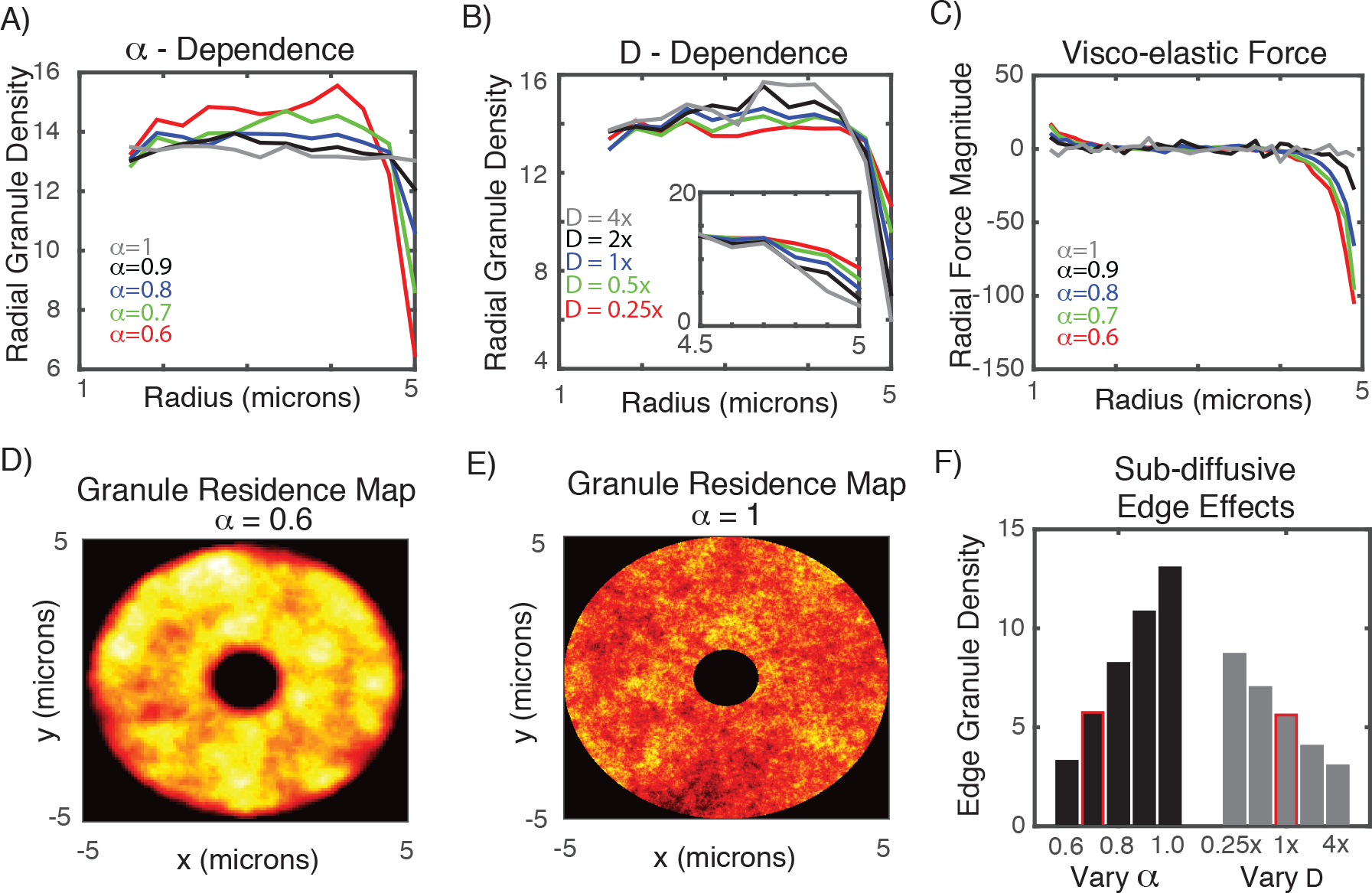
Influence of visco-elastic sub-diffusive effects on peripheral granule density. **Panel A:** Granule density as a function or radius for different values of the sub-diffusion exponent alpha. The rate of diffusion D is set to its base value here. **Panel B:** Radial granule density as a function of diffusion speed (D) for alpha=0.7. **Panel C:** Net radial visco-elastic force as a function of radius for different values of alpha, with D fixed at its base value. Negative values near the periphery indicate a net inward force. **Panels D, E:** Granule residency map showing the distribution of granules over time for two values of alpha (note the depletion zones near the inner and outer radii). **Panel F:** Effect of subdiffusive exponent (alpha) and diffusive strength (D) on the peripheral density of MTs. In the “Vary alpha” case, the D value corresponds to 1x (e.g. the base value). Similarly, for the “Vary D” case, alpha = 0.7. The highlighted bars most closely correspond to the calibrated D and “alpha” values used to model MT based diffusion for all other simulations to follow.

The explanation for this is subtle but readily explained by the basic assumptions of generalized Langevin dynamics. The physical mechanism often associated with visco-elastic sub-diffusion is that as a particle moves in a given direction, resistive forces on that particle build up due to interactions with the crowded, filamentous cellular environment; the more a particle moves in a given direction, the larger the resistive force becomes. If a particle is observed near a cell border, it is more likely that the particle was transported from more interior regions of the cell rather than more exterior regions. This would lead to an expected resistive force that would tend to move the particle back to the interior of the cell, introducing a bias not present in standard diffusion.

To confirm this explanation, we quantified the average radial component of the visco-elastic force as a function of radius within the simulated cell at steady state to generate a force (Fig. 4c). This force map quantifies the average, expected resistive force that a granule would be subject to as a function of radial location within the cell. When diffusion is close to normal (α=0.99), that force is effectively 0 everywhere. However, as diffusion becomes more anomalous, we begin to see a negative expected radial force near the cell border, suggesting a particle near the boundary would be expected to move inward rather than closer to the periphery. This is the source of the peripheral depletion of granules when motions are governed by visco-elastic sub-diffusion.

While this would be a general phenomenon in any system where visco-elastic sub-diffusion is present, it is specifically relevant here due to the dependence of this depletion effect on the speed of motion. The gray bars in Fig. 4f show that when the speed of motion is reduced by a factor of ¼, peripheral densities increase by roughly 50%. Interestingly, when MT’s are completely removed from beta cells via application of glucose + NOC, a roughly 1/4 - 1/3 reduction in *D* is observed (Figure 1c with data reproduced from (Zhu et al., 2015)) along with a roughly 50% increase in peripheral granule density.

We are not suggesting that this is the sole or even primary cause of the enrichment of peripheral granules near the periphery in response to MT removal. Glucose stimulation of Beta cells influences cells in a number of ways, including activating docking proteins that bind granules to the membrane. Additionally, the dynamics of MT’s significantly influence peripheral granule densities in other ways, independent of simply augmenting transport speed. None-the-less, it is expected based on this analysis that the net slow-down in motion would contribute to the peripheral enrichment observed experimentally when glucose and NOC are jointly applied to cells.

### Competition between membrane anchoring and MT binding regulates availability of peripheral granules

Here we consider how the dynamics of MT mediated motions influence granule localization and availability for membrane anchoring. A set of prior observations will allow us to assess what factors are important in understanding granule localization and constrain aspects of the computational model. In (Zhu et al., 2015), Zhu et al. quantified how granule density at the cell periphery changes when glucose, NOC, and glucose + NOC are applied to Beta cells. Briefly, they found that the application of either factor alone had relatively little influence on granule densities. However, when they were jointly applied, peripheral densities increased by on the order of 50%.

When initially studying peripheral granule accumulation without considering membrane anchoring, we found the model unable to account for these observations. We thus consider the joint effects of membrane anchoring and MT mediated transport, both of which are altered by glucose stimulation. To study how MT motion might influence membrane anchoring, we consider two possibilities for how granules anchor: 1) that any granule close enough to the periphery can anchor or 2) that only granules not bound to MTs can anchor. The latter possibility is motivated by the hypothesis that motions and forces subjected to granules by MT associated motors either prevent anchoring or substantially reduce anchoring affinity. Since we do not have anchoring protein affinity data, we consider the effects of low, medium, and high affinity (high not shown in data) as well as the absence of anchoring (relevant for NOC only treatment) on granule dynamics.

To understand the effect of MTs on the localization and availability of granules, we simulated the full model to steady state, performed both partial and complete removal of MTs (in different anchoring scenarios), and quantified the fold change in peripheral granule density. Results (Figure 5 b, c) show that in the absence of anchoring, neither partial nor complete removal of MTs alone has a significant effect on granule density and thus removal of MTs alone is not sufficient to explain granule enrichment when glucose + NOC is applied. The inclusion of anchoring can lead to the enrichment of peripheral densities, however those enrichment dynamics are only consistent with observations when MT unbound granules anchor to the membrane with low affinity. In this scenario, a roughly 40-50% increase in peripheral granules is found (Figure 5a), consistent with prior experimental observations (Zhu et al., 2015). When all granules can anchor independently of MT binding, enrichment occurs in the absence of any MT perturbation, contrary to observations. Alternatively, when affinity is too high, enrichment becomes extreme and once again independent of MT dynamics. In short, when anchoring is high affinity or all granules (MT bound and unbound) can anchor, MT properties have little effect on peripheral densities.

**Figure 5:**
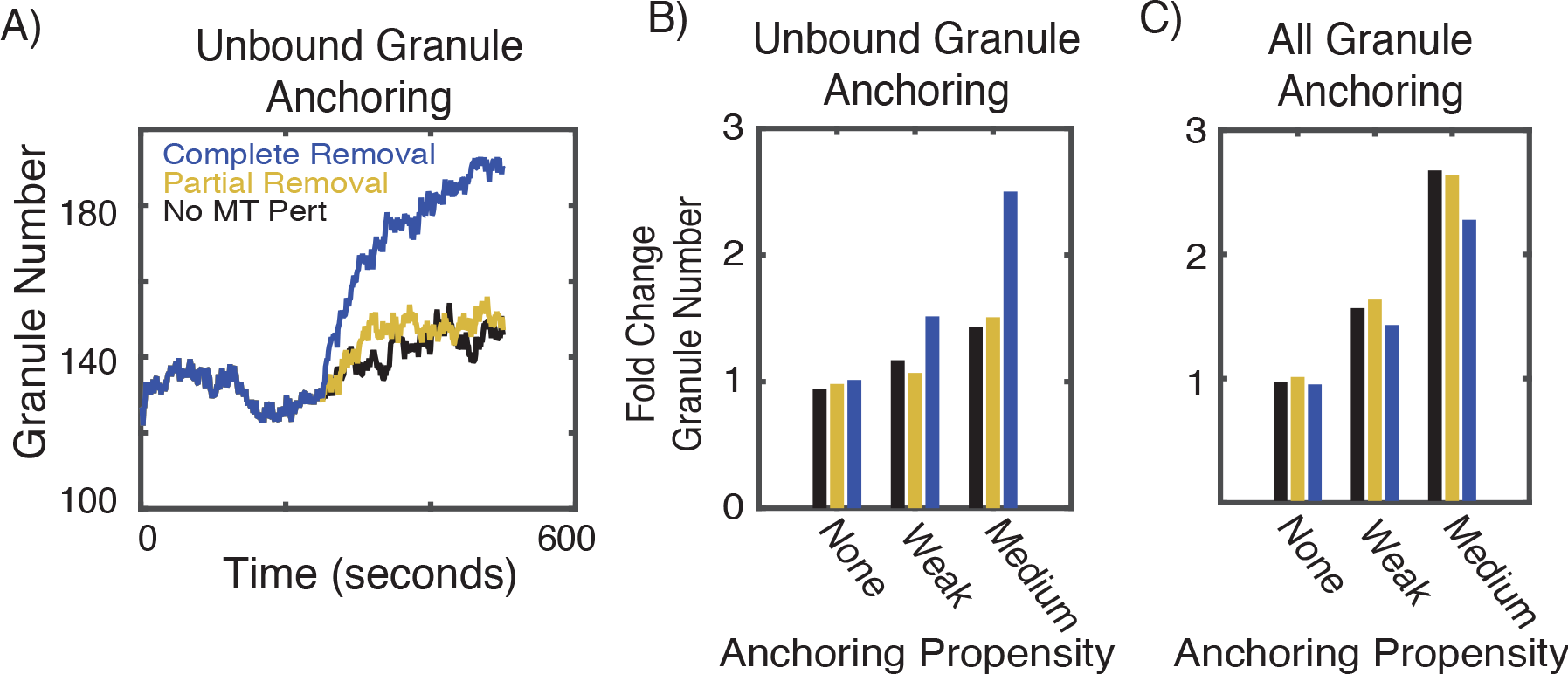
Influence of MT perturbations on peripheral granule density. **Panel A:** Number of granules within 250nm of the cell border as a function of time in simulations where only unbound granules are able to anchor to the membrane. This simulation shows the case for “weak” anchoring. The black curve indicates no MT perturbation is applied, red corresponds to removal of 1/3 of longer perturbations, and blue indicates complete removal of all MT’s (corresponding to Glucose + NOC treatment). **Panels B, C:** Quantification of the fold change in peripheral granule density after MTs are perturbed. Vertical axis shows the fold change in peripheral granule density after the relevant perturbation. The horizontal axis corresponds to different anchoring rates (none = 0, weak = 1/32, strong = 1/8). Slow MT dynamics are assumed so that the average lifetime is 1000 sec. Alternative simulations were performed with short MT lifetimes (10 sec, see SM Figure 2). In B, only MT unbound granules are allowed to anchor to the membrane. In C, all granules are assumed to be capable of anchoring.

We thus conclude anchoring is necessary to account for enrichment of peripheral granules upon glucose + NOC stimulation, but that only unbound granules should anchor and with low affinity. These results suggest MT’s may have a role in negatively regulating the availability of peripheral granules by binding them and making them unavailable for anchoring.

**Figure 6:**
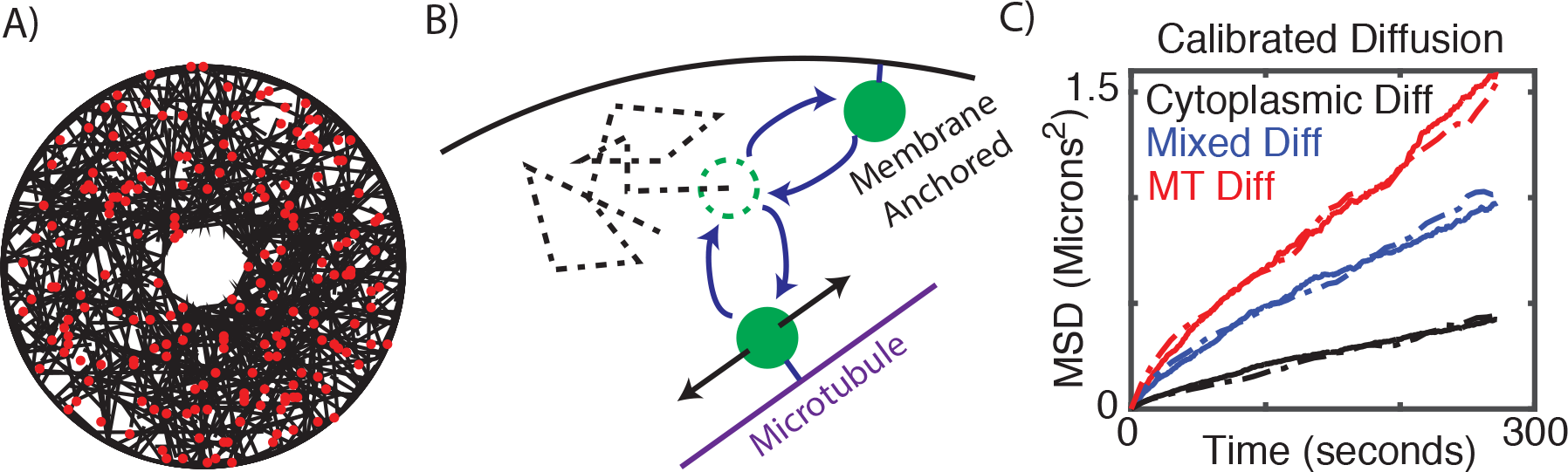
Description of model. **Panel A:** Snapshot of a simulation with 500 microtubules and 200 insulin granules. **Panel B:** Schematic of the basic model elements including free diffusion, binding of granules to MTs, diffusion along MT’s, and membrane anchoring. **Panel C:** Illustration of calibrated diffusion. Dashed curves are data and solid curves are simulated. Cytoplasmic and MT bound diffusion coefficients were calibrated based on the red and black curves and the relative fraction of time a granule spends in these two modes of motion was calibrated based on the blue curve.

## Discussion

Glucose homeostasis is tightly regulated at the systemic level, in both the amount of insulin in circulation and the response of peripheral tissues to insulin (including liver, skeletal muscles, and fat). This study here combines experimental test and modeling to investigate how beta cells regulate the amount of insulin to secrete at given stimuli. We focus on how the MTs in beta cells regulate the localization and anchoring of insulin granules to the plasma membrane, a pre-requisite for insulin secretion. Results here suggest that cytoskeletal factors contribute to tight regulation of insulin at the level of individual Beta cells.

While individual Beta cells can contain up to 13,000 individual insulin granules, only a few are secreted in response to glucose stimulation (Rorsman and Renstrom, 2003). Thus at the cellular level, significant negative regulation of GSIS must be present. A well-established key negative factor is the actin cytoskeleton, which ensures that only a small portion of vesicles are available to break the cell cortex and secreted (Wang and Thurmond, 2009). Here, we have identified two potential, alternative mechanisms by which MT dynamics contribute to this negative regulation. First, MTs near the cell periphery actively transport insulin granules away from the cell membrane. Second, traction forces generated by MT associated molecular motors prevent stable granule anchoring to the membrane, which is a precursor to exocytosis.

Both of these mechanisms are supported by prior observations. First, prior imaging (Zhu et al., 2015) demonstrated that depolymerization of the MT cytoskeleton substantially inhibited the removal of insulin granules from the membrane, supporting the conclusion here that MTs predominantly serve to remove granules from the cells surface. Secondly, recent work (Gandasi et al., 2018) demonstrated that membrane docking of granules is substantially inhibited in human type 2 diabetes. This along with the observation that MT density is increased in diabetic mouse models (Zhu et al., 2015) supports the conclusion that MT mediated transport prevents or inhibits anchoring of granules to the membrane.

Interestingly, both of these are consequences of an alteration in the MT structure near the cell membrane. Prior imaging has found the MT network in beta cells to be unusually unstructured and randomly oriented (Varadi et al., 2002; Zhu et al., 2015). However, results here demonstrate that in peripheral regions within ~250nm of the cell membrane, MTs are predominantly oriented parallel to the membrane. The two previously mentioned negative regulatory mechanisms are a direct consequence of this alteration in structure. As a result, the MT network acts like a sponge near the membrane that soaks up granules and transports them away from the periphery while preventing their membrane anchoring and stimulated release.

Co-alignment of MTs at the cell periphery can arise as a result of MT capture at the cortex or cell/cell junctions, which prevents MT catastrophe (Fukata et al., 2002; Gundersen, 2002; Schmoranzer et al., 2009; Stehbens et al., 2014; Watanabe et al., 2004; Zaoui et al., 2010) and thus can promote their turning by the actin retrograde flow (Bicek et al., 2009; Gupton et al., 2002) and polymerization along the cortex. Alternatively, subcellular signals localized at the plasma membrane, such as GSK3beta inactivation, can locally increase MT coating by MAPs, which, in turn, can promote excessive MT growth along the cell periphery (Kumar et al., 2009; Nishimura et al., 2012; Zhu et al., 2016). Such mechanisms that promote MT turning must also increase their lifetimes and stability, which is consistent with unusually high levels of stabilization that we observe in the peripheral MT bundles in beta cells. Thus there are a number of potential regulatory mechanisms under the control of cell differentiation and metabolic signals that could produce this aligned peripheral mesh and, as a result, tune insulin availability for GSIS.

Interestingly, the MT mediated withdrawal of peripheral granules does not require directed (i.e. ballistic) motor driven transport. That is, it is the topology of the MT network that influences cargo localization, not the specific motor dynamics (similar to (Ando et al., 2015)). Rather, the random granule motions observed in cells coupled with the structured nature of the network near the membrane is sufficient to generate directed motion of MT bound granules away from the membrane. It is interesting in this regard that while most studies concentrate on kinesin 1 as the main MT-dependent motor that transports insulin granules (McDonald et al., 2009; Varadi et al., 2002), the transport is likely driven by multiple motor transport involving both kinesin and dynein (Varadi et al., 2003). Furthermore, peripherally aligned MT arrays likely lack net polarity: there is no reason to anticipate that MTs growing along the cell periphery will be co-aligned. Furthermore, MT buckling at the periphery is capable of producing MTs with “reversed” polarity with their plus ends directed toward cell periphery (Zhu et al., 2016). In such a complex network, even solely plus-end directed molecular motors would promote non-directional transport.

The effects of MT configuration on granule distribution are predicted to persist under glucose stimulation conditions as well. MTs coaligned with the cell membrane and, accordingly, their functional consequences on granule dynamics are observed in both steady state (low glucose levels) and stimulated conditions (high glucose levels). Glucose stimulation does have two important consequences however. First, it leads to the activation of docking and exocytic machinery (Gandasi et al., 2018), which facilitates the secretion of those granules not interacting with the MT cytoskeleton. Secondly, it leads to depletion of stable long-lived MTs (Fig. 1 and (Zhu et al., 2015)) and replacing them by new, dynamic counterparts that are nucleated at the Golgi membrane (Zhu et al., 2015) and are characterized by rapid polymerization rates (Heaslip et al., 2014). While this does not lead to a gross reduction in MT density or readily detectable restructuring, it does reduce their lifetime, which, as our results suggest, lead to increased interaction between granules and the membrane and, subsequently, to increased secretion. Another way to interpret the result of increased MT dynamics is that it creates a pool of transiently “unbound” granules, which we show here to be the ones prone to accumulation at the cell periphery.

Here we used imaging and modeling to assess the consequences of MT dynamics specifically on secretion. However, much work is still needed to investigate how the biophysical properties of motors themselves as well as docking proteins influence secretion. One central hypothesis stemming from this work is that the motions and / or traction forces generated by the action of molecular motors on MTs parallel to the membrane inhibits stable membrane anchoring. Does this occur through the prevention of bond formation or the force dependent breaking of those bonds? Furthermore, how does the nature of the multiple motor transport these granules are likely subject to influence interactions with the membrane? Addressing these questions will require further experimental investigation of the biophysical interactions between the cytoskeleton, molecular motors, membrane docking proteins, and insulin granules.

These results do however suggest that there is a potential therapeutic merit in targeting the cytoskeleton to modulate Beta cell function. Recent imaging demonstrated that increased MT density was found to correlate with decreased secretion in mouse models (Zhu et al., 2015). While those results were correlative, our findings here indicate that *in silico* dense peripheral MT network interferes with the proper positioning of insulin granules for secretion. This result predicts that in fact the link observed in mouse models may be causal, and interference with MTs stability in beta cells might be used as an approach to increase insulin secretion efficiency. This idea is tempting because numerous MT-targeting small molecule compounds have been already considered or even used for cancer therapies. This potential has to be approached very carefully, given high toxicity of MT drugs on all cells and likely negative effects of prolonged MT destabilization on insulin biogenesis in beta cells specifically. Nevertheless, one can envision that, in the future, locally delivered and released in a time-restricted manner MT destabilizers could be applied to facilitate insulin secretion and overcome hyperinsulinemia in patients. If proposing such an intervention is too bold, it is more realistic that future studies will identify specific MT stabilizing MAPs, which are responsible for high MT density in diabetes models. Then, potential therapies would become possible that specifically target these MT-binding proteins.

## Author Contributions

K.M.B. performed imaging and experimental data analysis. K.-H.H. provided samples and performed imaging. D.Y. designed image analysis algorithm and program. G.G. supervised mouse work and edited the manuscript. I.K. supervised the experimental part of the study, provided conceptual insight, and edited the manuscript. W.R.H. designed computational simulations, supervised the computational part of the study, provided conceptual insight, and wrote the manuscript. There is no conflict of interest for all authors.

### Acknowledgements

This work was supported by a National Science Foundation grant DMS1562078 (to WRH), National Institutes of Health (NIH) grants R35-GM127098 and (to I.K.) and R01-DK106228 (to I.K. and G.G.). K.M.B. was supported by an NIH training grant R25-GM062459 “Initiative for Maximize Student Diversity” (IMSD) (Sealy, PI). KH was supported by Lilly Innovation Fellowship Award (LIFA) UNIV59676. We utilized the core(s) of the Vanderbilt Diabetes Research and Training Center (funded by NIH grant DK020593). We thank Hamida Ahmed for technical help.

## Materials and Methods

### Discrete Model Description

This discrete model accounts for four essential features that impact the transport of granules: 1) transport along MTs, 2) transport independent of MTs, 3) binding and unbinding of granules to MTs, and 4) tethering of granules (that are sufficiently close) to the cell membrane, which renders them immobile. We briefly discuss how each of these features is encoded into the model and how aspects of it are calibrated to data.

### Modeling MT independent transport

We model the cell as a 5 μm radius circle with a 1 μm hole cut out (representing the nucleus). A regular 2D grid is constructed on this cellular domain and motion of granules on this lattice is modeled as a sub-diffusive random walk. Granule motion is assumed to obey the equation of motion for over-damped Fractional Langevin Equation (FLE) representing viscoelastic sub-diffusion

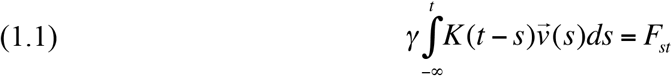

where *v* is the velocity of the granule, *γ*is the drag coefficient for the granule, ***K*** is a memory kernel encoding viscoelastic effects, and *F_st_* is the stochastic forcing obeying the appropriate fluctuation dissipation theorem (Kubo). Here the drag coefficient satisfies *γ*=2*k_B_T*/(Γ(1+α)*D*) where *k_B_T* is the standard thermal noise constant and *D, α* are estimated from data (see (Lutz, 2001) for further details of the FLE). The kinetic lattice Monte Carlo method for simulating the over-damped FLE from (Fritsch and Langowski, 2012) is used to simulate motion.

### Constructing the MT network and modeling MT mediated transport

To simulate MT mediated motion, we must first populate the *in silico* cell with a microtubule network. We do so by essentially growing a network of five hundred discrete, independent, and non-interacting microtubules. Of these, 250 are short (mean length 2 microns) and 250 long (mean 5 microns). For each MT, we specify a start point and initial growth direction. The MT is then grown in a strait line until it interacts with the cell periphery (if it does so at all). If the growing MT interacts with the cell border, it either terminates with probability *p* or bends and grows parallel to the cell periphery with probability *1-p*. For most simulations a value of *p=0* is chosen, corresponding to a peripherally aligned population of MTs forming. However in Figure 2, the effect of this MT structure parameter is considered. For simplicity, we assume the MTs are fixed in place once grown, and thus do not model the detailed dynamics of remodeling of the MT cytoskeleton by motors themselves (Hillen et al., 2017; White et al., 2015). The model does incorporate dynamic instability (Goodson and Jonasson, 2018) of MT’s through removal and replacement of MT’s with a specified rate. We fix the average MT lifetime at 1000 seconds, though do also consider the effects of short MT lifetimes (10 sec, SM Figure 2). As removal / replacement of MTs has much the same effect on granule motility as MT binding / unbinding, which is considered in somewhat more detail, we do not exhaustively explore the effect of MT catastrophe dynamics.

Motion of granules is strongly sub-diffusive even in the absence of actin, suggesting that individual MT motors are not moving granules along in a directed fashion along them (this would be super-diffusion). Furthermore, each granule likely has a number of motors bound to it at any given time that are constantly competing to be the driver of motility. We thus do not model the dynamics of individual motors. Rather, MT mediated granule motion is modeled as a 1D sub-diffusive random walk on the MT to which the granule is bound. The same FLE equation of motion (Equ. 1.1 above) describes this motion and the same method of simulation is used.

### Granule binding, unbinding, and tethering

It is highly likely that granules bind and unbind from the MT lattice and thus their aggregate motion is in some sense an interpolation of motion in those two forms. We assume that an unbound granule can bind to any MT within 250 nm of its center (assuming a 150nm granule radius along with an additional ~100nm reach of the motor head) with a per MT rate of binding k_on_. Similarly, granules are assumed to unbind from MT’s with a rate of k_off_. These two rate constants are not individually accessible as we don’t know what specific motors are involved or how granules interact with the dense network of MTs. As discussed shortly however, we can calibrate their ratio based on data to ensure that relative fraction of time each granule spends undergoing (un)bound motility is appropriate. We thus fix the rate k_off_ =1/30 to represent a roughly 30 second bound lifetime, which is in the same range found for the kinesin Kip2 (Hibbel et al., 2015), and calibrate k_on_ accordingly. Switching of granules from one MT to another is not directly modeled. However when a granule unbinds from a MT, it can re-bind to any other nearby MT. All simulations were carried out with k_off_ =1/15, 1/60, yielding similar results.

Anchoring of granules to the cell membrane is modeled similarly with granules within 250nm of the cell border binding with a rate constant k_T_ and unbinding with a rate k_U_. We again do not have estimates for these rate constants or any way to constrain them directly. In this study however, we vary the affinity by fixing the unbinding rate and modulating the magnitude of k_T_ to determine how the relative strength of anchoring, or in other words, the relative fraction of time an unbound granule spends tethered to the membrane, influences dynamics (see Figure 3). We further consider two possibilities for which granules can / cannot tether: 1) that all granules can tether or 2) that only unbound granules can tether.

### Calibrating motility and binding/unbinding parameters

While all parameters of this model are not estimable, some are. In particular, using granule motility data we can extract the values of the sub-diffusive exponent (α), the diffusion coefficients for bound and unbound granule motion (*D* for each), and the relative fraction of time the granule spends in the bound and unbound states.

The data we use for this is derived from (Zhu et al., 2015) where mean squared displacement (MSD) of granule motion was measured in control cells, cells with the MT cytoskeleton removed, and cells with the actin cytoskeleton removed. First, from (Tabei et al., 2013) it was determined that α=0.75. To calibrate the diffusion constants, we will rely on MSD data. For simplicity, we will assume that in the absence of actin, all motion is MT mediated, and in the absence of MT’s, all motion is actin mediated. We can thus use the data where actin is removed to calibrate the diffusion constant for MT motion, and the data where MT’s are removed to calibrate the diffusion constant for non-MT motion. See Figure 1c (black and red curves) for calibration results.

We can additionally use the control MSD data to calibrate the relative fraction of time each granule spends in bound and unbound states. The idea here is that the more time a granule spends in the MT bound state, the higher its MSD will be and vice versa. We fix the value of k_off_ and vary the value of k_on_ until the MSD data of the amalgamated diffusion matches that of control data (blue curve in Figure 1c). For a list of all parameters for the discrete model, see Table 1.

**Table 1:** Synopsis of the discrete model parameters.

| Parameter Name | Value | Units |
| --- | --- | --- |
| Number of MTs | 500 | Number |
| Number of Granules | 1000 | Number |
| Cell Radius | 5 | $\mu\text{m}$ |
| Nucleus Radius | 1 | $\mu\text{m}$ |
| Granule Radius | 0.3 | $\mu\text{m}$ |
| Sub-diffusion Exponent ($\alpha$) | 0.75 | None |
| MT Bound Diffusion Constant ($D_1$) | 0.015 | $\mu\text{m}^2 / \text{sec}$ |
| 2D Grid Diffusion Constant ($D_2$) | 0.005 | $\mu\text{m}^2 / \text{sec}$ |
| MT Mean Length (short, long) | 2, 5 | $\mu\text{m}$ |
| MT Length Standard Deviation | 2 | $\mu\text{m}$ |
| MT Catastrophe Rate | 1/1000 | 1 / sec |
| Granule / MT binding rate ($k_{\text{on}}$) | 1/8 | 1 / (sec*MT) |
| Granule / MT unbinding rate ($k_{\text{off}}$) | 1/30 | 1 / sec |
| Granule / MT binding radius | 0.25 | $\mu\text{m}$ |
| Granule Tethering rate ($k_T$) | 0 | 1 / sec |
| Granule Untethering rate | 1/30 | 1 / sec |
| Granule Tethering Radius | 0.25 | $\mu\text{m}$ |

### Simulation protocol

All simulations begin with each of the 1000 granules randomly placed in the cell for a pseudo-uniform distribution. The time step for simulations granule motions is chosen to be ΔT=10ms with simulations running nominally for 300 sec to achieve a steady state. The kinetic lattice Monte Carlo method requires that the spatial grid size be chosen appropriately so that Δ*x* = (2*D* Δ*T* ^*α*^)^0.5^. Since the diffusion constant for the MT and non-MT mediated motions are different, this yields spatial step sizes of 18nm for the 2D lattice motions and 30nm for the 1D motions on MTs. In select simulations, decreasing the time step to ΔT=5ms did not alter results. In all results presented, averages of 50 independent simulations are presented unless otherwise stated.

### Mice

Mouse usage and handling followed the protocol approved by the Vanderbilt Institutional Animal Care and Usage Committee for Dr. Gu. Wild type CD-1 (ICR) mice were purchased from Charles River Inc. (Wilmington, MA). All mice were bred and handled following protocols approved by the Vanderbilt Institutional Animal Care and Use Committee (IACUC). All mice used were 8-10 weeks of age.

### Islet isolation

Islets isolation followed the previously described procedure (Brissova et al., 2002). Briefly, mouse pancreata were distended by injecting 3 mL 0.8 mg/mL collagenase P (Sigma, St Louis, MO) through the bile duct and digested at 37°C for 20 minutes. Islets were hand-picked and cultured to recover in Gibco™ RPMI 1640 Medium (Thermo Fisher, Waltham, MA) containing 11mM glucose, 10% heat inactivated FBS (Atlanta Biologicals, Flowery Branch, GA), 100 IU/mL penicillin, and 100 µg/mL streptomycin.

### Immunofluorescence

Isolated mouse Islets were treated with 2.8 mM (low) or 20 mM (high) glucose in RPMI media for two hours, and fixed with 4% paraformaldehyde in PBS with 0.1% saponin (Sigma, St Louis, MO). Immunofluorescence followed the described procedure (Zhu et al., 2015). Briefly, fixed islets were stained with primary antibodies at 4 °C for overnight followed by another staining with fluorophore-conjugated secondary antibodies. After each staining, islets were washed using PBS with 0.1% saponin for three times. After staining, islets were mounted with Vectashield mounting media (Vector Labs, Burlingame, CA) for microscopy. Primary antibodies used are rabbit anti-β-tubulin (Abcam, Cambridge, MA), guinea pig anti-insulin (DAKO, Houston, TX), rabbit anti-detyrosinated tubulin (Millipore, Burlington, MA), and mouse anti-E-Cadherin (BD Biosciences, San Jose, CA). Secondary antibodies used are goat anti-rabbit IgG-Alexa Fluor 488 (Abcam, Cambridge, MA), goat anti-mouse IgG-Alexa Fluor 488 (Invitrogen, Grand Island, NY), and goat anti-guinea pig IgG-Alexa Fluor 650 (Thermo Fisher, Waltham, MA)

### Microscopy and Image processing

All images were captured using Nikon Eclipse A1R laser scanning confocal microscope equipped with a CFI Apochromat TIRF 100X/1.45 oil objective. The microscope is driven by Nikon Elements software. For directionality analysis, oversampled image stacks (50nm^3^ voxeles) were acquired and thereafter deconvolved by NIS Elements Software using Richardson and Lucy algorithm (15 iterations). All images presented in figures were single-slice confocal images, where the brightness and contrast were adjusted consistently across every image to better present small structural features.

### Image Analysis

Image analysis algorithm was developed to determine the alignment of neighborhood structure at internal point within tubulin images with that at nearest point on the border. Beta cells within an islet were selected based on their ability to express insulin. Single slices from a deconvoluted confocal stack were used for analysis. Taken into consideration that MT width is below the resolution limit of microscopy, neighborhood block size was approximated to the pixel size of the oversampled confocal image (50nm^2^). Analysis was applied within a mask based on thresholded tubulin images. The local orientation at each pixel of tubulin image was derived using method described in (Feng and Milanfar, 2002). Cell outline curve, manually constructed based on the E-cadherin staining, was smoothed and used to estimate orientation of cell border. Each pixel in tubulin image was associated with a pixel of the boundary curve, nearest to it. Per pixel not excluded by cell and tubulin threshold masks, angle difference between local orientation and orientation of boundary at nearest pixel was calculated, zero indicating perfect parallel alignment. Results were weighed according to variance of local orientation, to avoid data from lumps of tubulin bands of excessive density and MT crossings with ambiguous configuration in the results. All pixels were manually sorted according to their distance from the cell boarder into bins of 0.5 µm.

---

**SM Figure S1:**
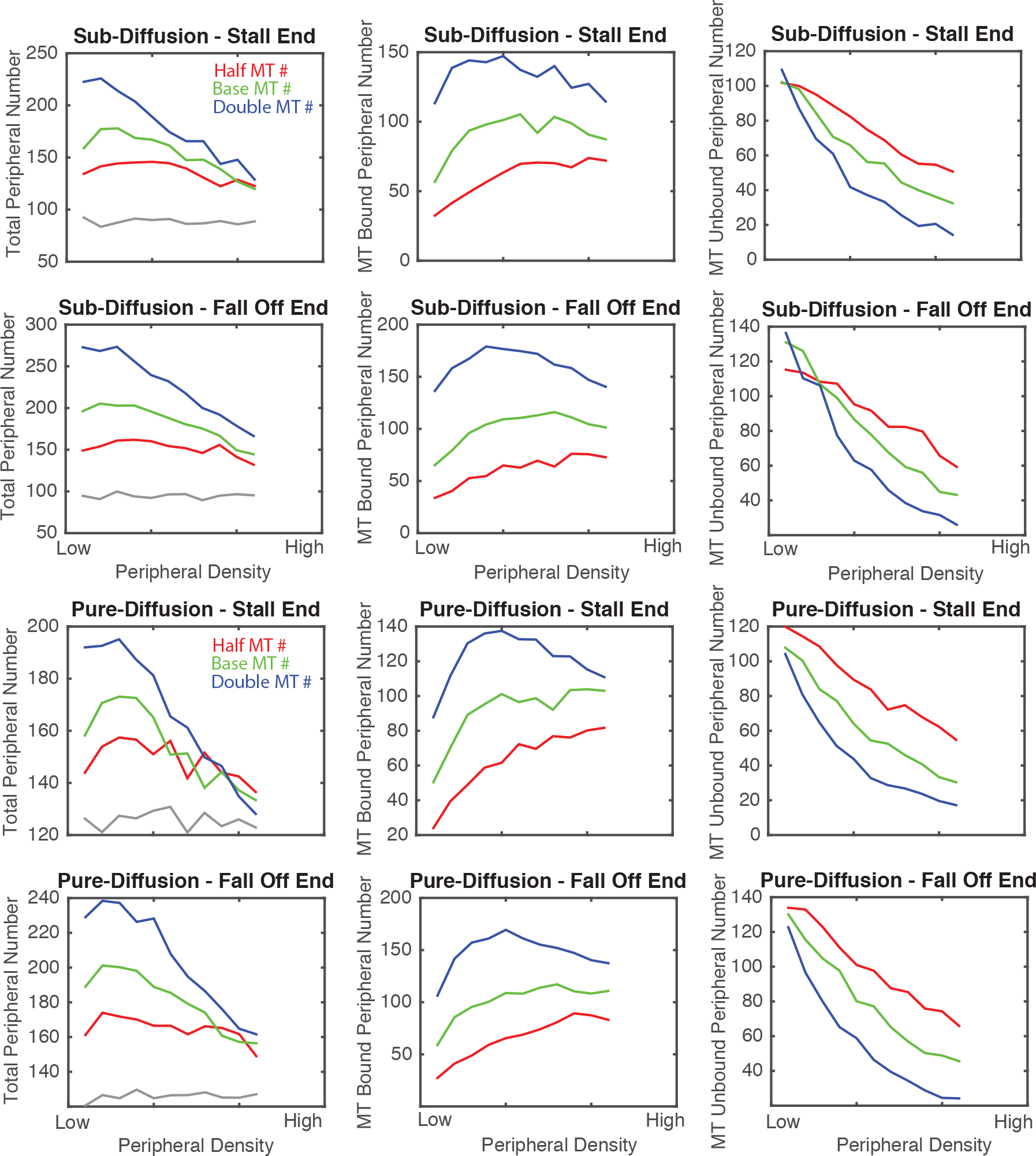
Role of motor stalling and type of diffusion on peripheral granule density. Figure 2 A-C assessed the effect of MT dynamics on the peripheral localization of MT bound, unbound, and total insulin granules. The top row here is a reproduction of that data for comparison purposes. The second row shows a similar set of simulations where instead of assuming motors stall when reaching the end of MT, they walk off the end and the granule dis-associates from that MT. The third and fourth rows show similar simulations where granule dynamics are governed by standard diffusion rather than sub-diffusion. These results show that the type of motion (standard versus sub diffusion) and the assumptions about the dynamics of motors at the end of MTs have little effect on results of this study.

**SM Figure S2:**
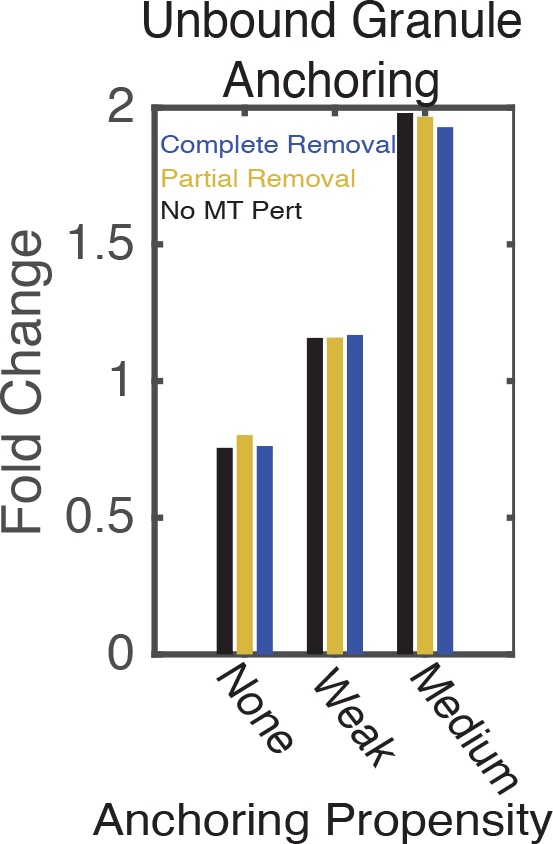
Impact of MT lifetime granule density after MT perturbation. Quantification of the fold change in peripheral granule density after MTs are perturbed when MT lifetimes are short (10 sec, compared to 1000 sec in all prior simulations). All plotting conventions are the same as Figure 5B. The horizontal axis corresponds to different anchoring affinities (none = 0, weak = 1/32, strong = 1/8). Results show that if MT lifetimes are short, MT removal has little effect on peripheral granule density.

